# Evaluating the Reliability of AlphaFold 2 for Unknown Complex Structures with Deep Learning

**DOI:** 10.1101/2022.07.08.499384

**Authors:** Hao Xiong, Long Han, Yue Wang, Pengxin Chai

**Affiliations:** Department of Molecular Biophysics and Biochemistry, Yale University, New Haven, The United States

## Abstract

Recently released AlphaFold 2 shows a high accuracy when predicting most of the well-structured single protein chains, and subsequent works have also shown that providing pseudo-multimer inputs to the single-chain AlphaFold 2 can predict complex interactions among which the accuracy of predicted complexes can be easily determined by ground truth structures. However, for unknown complex structures without homologs, how to evaluate the reliability of the predicted structures remains a major challenge. Here, we have developed AlphaFold-Eva, a deep learning-based method that learns geometry information from complex structures to evaluate AlphaFold 2. Using different types of sub-complexes of the central apparatus and recently released PDB data, we demonstrate that the reliability of unknown complex structures predicted by AlphaFold 2 is significantly affected by surface ratio, contact surface and dimension ratio. Our findings suggest that the reliability of predicted structures can be directly learned from the intrinsic structural information itself. Therefore, AlphaFold-Eva provides a promising solution to quantitatively validate the predicted structures of unknown complexes, even without a reference.

## Introduction

The prediction of protein structure from amino acid sequence information alone has been a century problem. In the 14^th^ CASP (Critical Assessment of Structure Prediction) meeting, DeepMind introduced an updated version of the protein structure prediction model, AlphaFold 2, which showed much higher performance than its previous version (AlphaFold) (Jumper et al. 2021). The average global distance test score of AlphaFold 2 could reach over 85, while AlphaFold was only about 60 (Callaway et al. 2020). The main technique breakthrough in AlphaFold 2 is that it changes the neural network architecture from convolutions to attention architecture. In this attention architecture, it makes the information exchange between multiple sequence alignment (MSA) and protein structure possible (Jumper et al. 2021).

Although AlphaFold 2 can provide such high accuracy, the whole architecture is trained on individual protein chains, including many proteins whose structure is solved in complex with other proteins (Evans et al. 2021). Therefore, whether it can predict complexes is still not well understood. Recently, several unpublished works and personal trials have been shown that providing pseudo-multimer inputs (e.g. chains joined with a flexible linker) is a way to test the accuracy of AlphaFold 2 on complexes (Baek et al. 2021). Moriwaki and Ohue showed that AlphaFold 2 may predict heterocomplexes by adding a long linker to connect the two sequences (Moriwaki et al. 2021). Ko et al. (2021) used AlphaFold 2 to predict protein-peptide complex and concluded that for about 40% of cases, the peptide structures can be modeled with good accuracy. Bryant et al. (2021) applied AlphaFold 2 to predict the structure of heterodimeric protein complexes and stated that using the default AlphaFold 2 protocol, 44% of the models in the test set can be predicted accurately. Meanwhile, several AlphaFold-based algorithms have been developed. For example, AlphaFold-linker adds a 21-residue repeated Glycine-Glycine-Serine linker between each chain to predict the complexes. AlphaFold-Gab runs the AlphaFold with a 200-residue gap in the residue index between chains (Ovchinnikov et al. 2021). Ghani et al. (2021) improved the docking of the protein model by a combination of AlphaFold 2 and ClusPro.

These works have shown the high potential performance of the original trained AlphaFold 2 model on complexes but also leave some open questions. First, how accurate AlphaFold 2 is on complex structures is still not thoroughly investigated. Second, when dealing with complex structures, what factors will influence the accuracy of AlphaFold 2? More importantly, all recent works judge the reliability of AlphaFold 2 on complex structures by comparing them with the ground truth structures. What if there are no ground truth structures (actually, this is the most common case when we explore the unknown structures), how can we evaluate the reliability of AlphaFold 2 in this situation? The answers to these questions form the interest of this work.

Here, we use the default AlphaFold 2 to predict four kinds of protein-protein complexes (homodimer, homotrimer, heterodimer, and heterotrimer) to understand the accuracy of AlphaFold 2 on complexes. Then we analyze nine factors (molecular weight, sequence length, largest and smallest dimension sizes, dimension ratio, complex total volume, total surface, contact surface, and surface ratio) that might influence the accuracy of AlphaFold 2 on complexes. Random forest is used to do feature-importance selection (Wu et al. 2019) and finally AlphaFold-Eva, a deep learning-based method, is trained to evaluate the reliability of the AlphaFold 2 on complexes without ground truth structures.

## Methods

In this work, we prepare three benchmark sets of protein-protein complexes. Since the cut- off date for training AlphaFold 2 is 2020-5-14, the first set consists of 469 protein-protein complexes randomly selected from the RCSB PDB database where the released date is before 2020-5-14. The second set consists of 444 protein-protein complexes where the released date is after 2020-5-4. The third set is from our unpublished (Central Pair Complexes) and recently released PDB files. The first two datasets are used to testify the accuracy of AlphaFold 2 on complexes. Then some important features are extracted from the complexes and used to train the AlphaFold-Eva model, and the third datasets are applied to test the accuracy of AlphaFold-Eva (please see details later). Note that the source code of AlphaFold 2 was retrieved from the GitHub repository and run locally to guarantee there is no updated training database of AlphaFold 2.

We evaluate the accuracy of protein-protein complexes using RMSD_All pairs because it is used as one of the assessments of how well a submitted structure matches the known in CASP (Damm et al. 2006). We treat RMSD_All pairs below 5 Å as good predictions and above as bad predictions following the protocol of TM-score (Xu et al. 2010). The build-in package Chain-Paring of Chimera X is used to calculate the RMSD_All pairs between the ground truth structures and predicted ones (Pettersen et al. 2021). It should be noted that RMSD is calculated by local similarity, that is, it will only calculate one chain and ignore other chains when dealing with complexes. To solve this problem, the package *Coot* is used to link all the complex chains together without modifying the structure information (Emsley et al. 2004). After this processing, the calculated RMSD is for global similarity. UCSF Chimera X is used to visualize complex structures (Pettersen et al. 2021).

## Results and Discussion

### Accuracy of AlphaFold 2

The total performance of AlphaFold 2 is shown in Figure 1 where the accuracies of homomer and heteromer are 61.7% and 48.8%, respectively. Labels 0 and 1 mean good and bad predictions separately. Figure 1 indicates the performance of AlphaFold 2 is better for homomer than that for heteromer. This result is consistent with Evans et al. (2021) where they guessed in the homomeric cases the MSA (multiple sequence alignment) is easier to encode evolutionary information than that in heteromeric interfaces.

**Figure 1.**
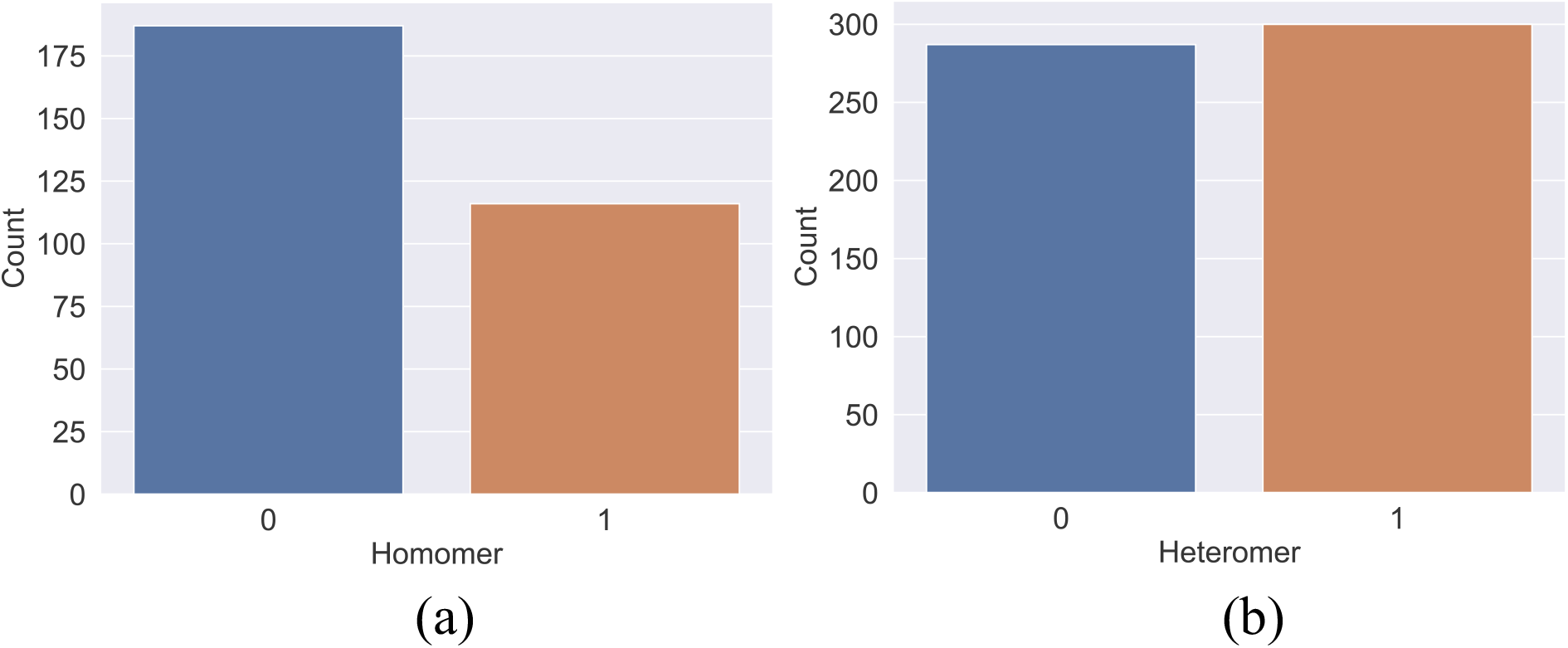
The performance of AlphaFold 2. (a) homomer; (b) heteromer datasets. Labels 0 and 1 mean good and bad predictions respectively. The threshold value of RMSD is 5 Å. The total performances of AlphaFold 2 in homomer and heteromer are 61.7% and 48.8%, respectively which is consistent with Evans et al. (2021).

### Feature Importance

In this section, we discuss nine features that might influence the accuracy of AlphaFold 2, that is, molecular weight, sequence length, largest and smallest dimension sizes, dimension ratio, complex total volume, total surface, contact surface, and surface ratio. The reason we choose these nine features is that all of them can be easily obtained from the RCSB PDB database and Chimera X (Pettersen et al. 2021).

Figure 2 shows the definition of the contact surface and surface ratio where *S*_*1*_, *S*_*2*_, and *S*_*3*_ are the surface of proteins 1, 2 and 3 separately. *S*_*c1*_ is the contact surface between proteins 1 and 2. Therefore, the surface ratio between proteins 1 and 2 is 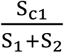 as shown in Figure 2(a). Following this logic, we can get the surface ratio in homotrimer (Figure 2b) and heterotrimer (Figure 2c). The structure of dimension size is determined by Chimera X, and the largest and smallest dimension sizes are treated as two input features. The dimension ratio is equal to 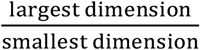. The rest of the features can be understood easily and are not explained here.

**Figure 2.**
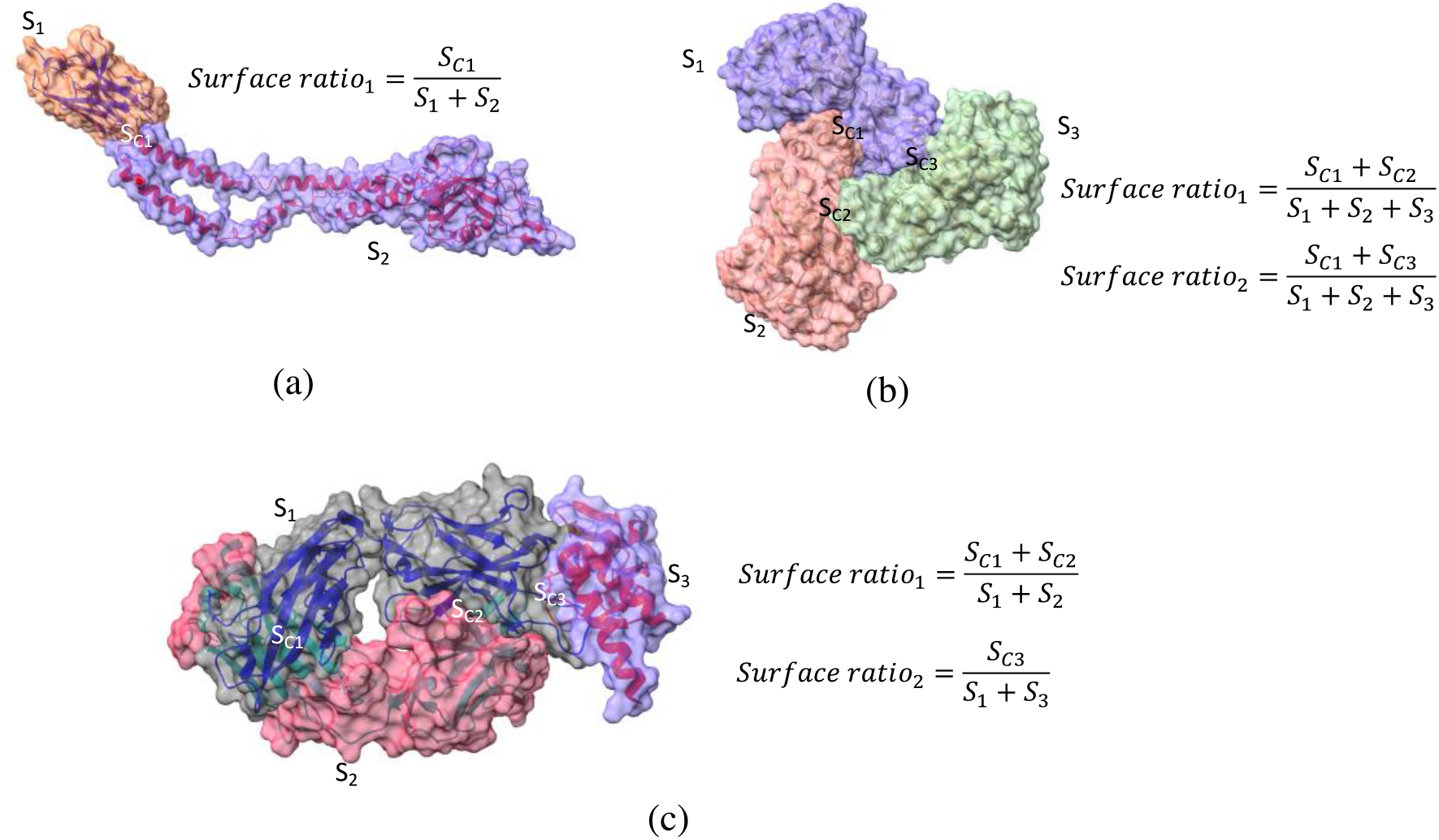
Definition of the contact surface and surface ratio in heteromer and homomer. S1 is the surface of protein 1, S2 is the surface of protein 2, S3 is the surface of protein 3, and Sc1 is the contact surface between proteins 1 and 2. the surface ratio between proteins 1 and 2 is 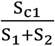. Following this logic, we can get the surface ratio in homotrimer (Figure 2b) and heterotrimer (Figure 2c).

Random forest is used to do feature selection because it provides good predictive performance, low overfitting, and easy interpretability (Wu et al. 2019). Training features are the nine features mentioned above, and the training target is AlphaFold 2 prediction results (0 and 1). Before applying random forest, MinMaxScaler is used to transform all the features into the range [0,1] (Ioffe et al. 2015). The result of feature selection is shown in Figure 3 (a). It indicates that surface ratio, contact surface, and dimension ratio are the top 3 factors that influence the accuracy of AlphaFold 2. Figure 3 also demonstrates that molecular weight and sequence length show small impacts on AlphaFold 2 prediction, in consistence with Ko et al. (2021) work where they stated that modeling accuracy does not depend on the sequence length of complexes.

**Figure 3.**
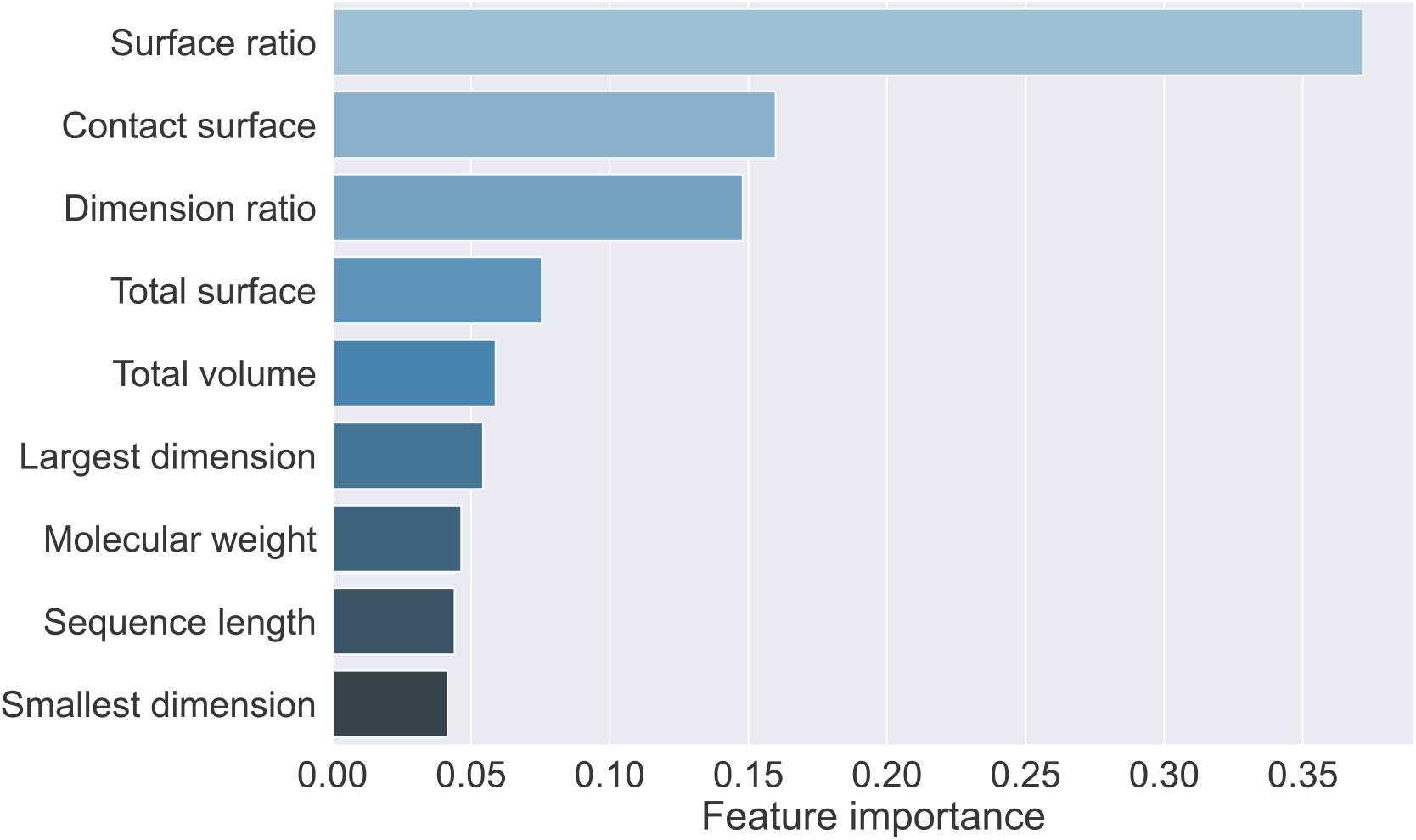
Feature selection using random forest. Training features are the nine features mentioned above, and the training target is AlphaFold 2 prediction results (0 and 1). Surface ratio, contact surface, and dimension ratio are the top 3 factors that influence the accuracy of AlphaFold 2. It also demonstrates that molecular weight and sequence length show small impacts on AlphaFold 2 prediction, in consistence with Ko et al.’s (2021) work.

Next, we further analyze the top 3 factors in our dataset as shown in Figure 4. Figure 4 (a-b) shows that the good prediction (label 0) usually has a higher surface ratio and contact surface than the bad prediction (label 1), demonstrating that with a large contact surface (or surface ratio), AlphaFold 2 can extract more information from the interaction surface to increase the prediction accuracy of complexes. Figure 4 (c) shows the distribution of surface ratio in heteromer and homomer where homomer complexes usually have a higher surface ratio than heteromer ones, indicating that AlphaFold 2 can yield a better prediction in homomer complexes, which agrees with our (Figure 1) and Evans et al.’s (2021) predictions. Figure 4 (d) shows the distribution of dimension ratio in all structures, indicating that in good predictions, the value of dimension ratio is usually less than that in bad predictions. In other words, AlphaFold 2 shows better performance on more cubic-shaped structures than rectangular ones.

**Figure 4.**
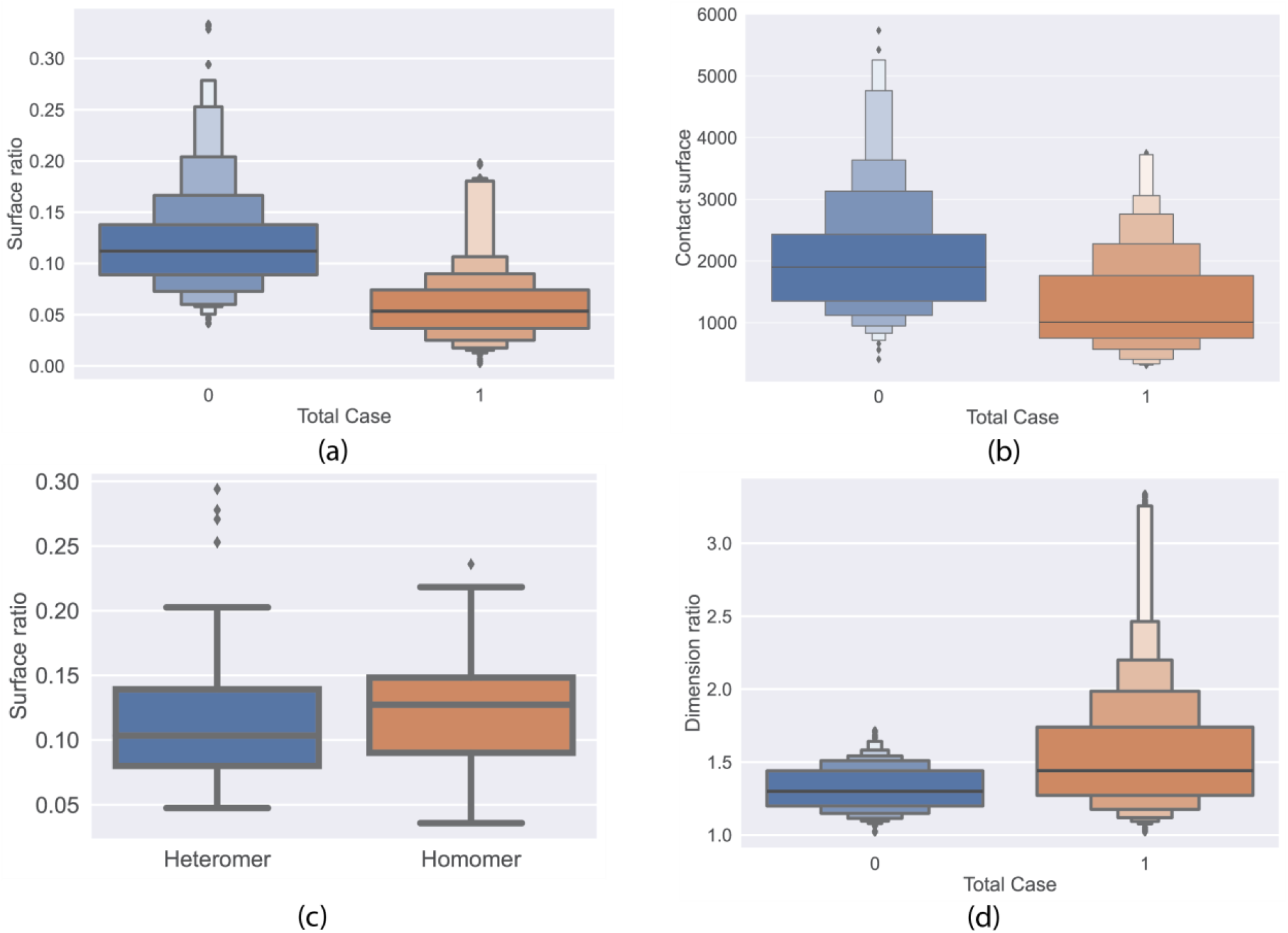
Analysis of the top 3 features. (a) Distribution of surface ratio in all structures, where labels 0 and 1 mean good and bad prediction respectively. (b) Distribution of contact surface in all structures; (c) Distribution of surface ratio in heteromer and homomer. Note that we only analyze the good predictions. It indicates that AlphaFold 2 yields a better prediction in homomer than heteromer because homomer complexes usually have a larger surface ratio than heteromer ones, in consistence with our prediction (Figure 1) and Evans et al.’s (2021) results (d) Distribution of dimension ratio in all structures.

Carrying with this knowledge, we analyze the AlphaFold 2 prediction results under some specific conditions as shown in Figure 5, where threshold values of dimension and surface ratios are 1.5 and 0.1 separately based on Figure 4 (a) and Figure 4 (d). Figure 5 (a-b) presents that when the dimension ratio of the complex is larger than 1.5, the accuracy of AlphaFold 2 is about 24%. However, when the dimension ratio is less than 1.5, the accuracy of AlphaFold 2 increases to about 59%. Figure 5 (c-d) demonstrates that when the surface ratio is less than 0.1, the accuracy of AlphaFold 2 is about 25%. However, when the surface ratio is larger than 0.1, the accuracy of AlphaFold 2 increases to about 90%. Therefore, we treat the dimension ratio less than 1.5 and surface ratio larger than 0.1 as good conditions (features) and vice versa. Meanwhile, this sudden change of accuracy also demonstrates that surface ratio (changing from 25% to 90%) has more impact than dimension ratio (changing from 24% to 59%), in consistency with our conclusion in Figure 3.

**Figure 5.**
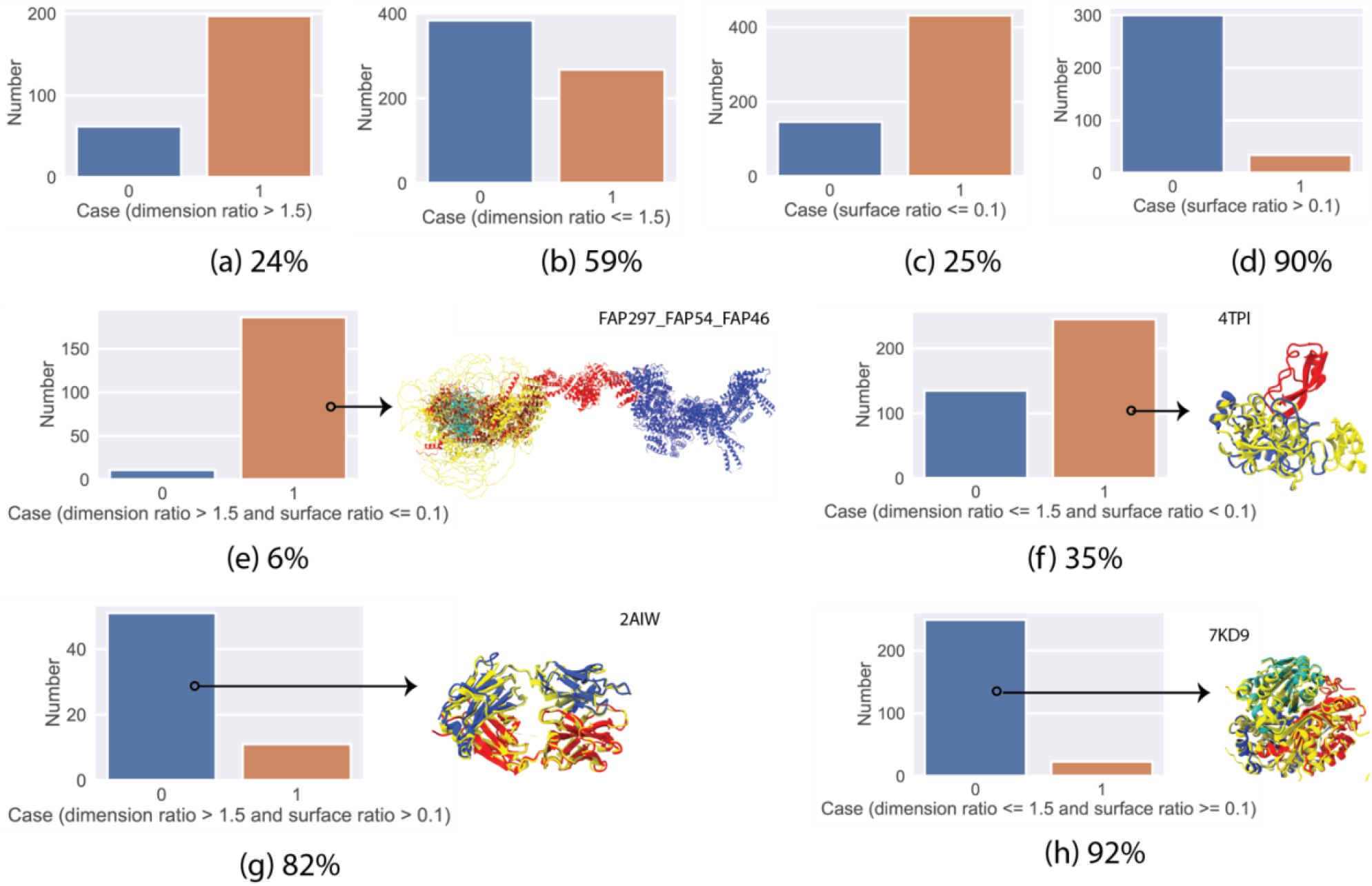
AlphaFold 2 prediction results under some specific conditions. (a) Dimension ratio >1.5; (b) Dimension ratio <=1.5; (c) Surface ratio <=0.1; (d) Surface ratio >0.1; (e) Dimension ratio >1.5 and surface ratio <=0.1; (f) Dimension ratio <=1.5 and surface ratio <0.1; (g) Dimension ratio >1.5 and surface ratio > 0.1; (h) Dimension ratio <=1.5 and surface ratio >=0.1. The structures in panels (e-h) are some typical ones in its correponding conditions and visualized are the ground truth structures (colored by chain) and predicted structures (yellow).

Figure 5 (e) shows the AlphaFold 2 prediction results with two bad conditions (dimension ratio >1.5 and surface ratio <0.1). In this case, it is not a wise choice to use AlphaFold 2 to predict the complexes where the accuracy is about 6%. The mess of AlphaFold 2 prediction (FAP297_FAP54_FAP46, Figure 5e) confirms our thought where the ground truth structures are colored by chain and predicted structures yellow. In contrast, if the complexes contain two good conditions (dimension ratio <1.5 and surface ratio >0.1) as shown in Figure 5 (h), AlphaFold 2 can yield the best predictions (i.e., 7KD9) with an accuracy of 92%. At the intermediate (one good and one bad), the accuracies of AlphaFold are 35% (Figure 5f) and 82% (Figure 5g), respectively.

### Neural Network

For the complex structures we know, it is easy to determine whether the prediction of AlphaFold 2 is good or not. However, for the unknown complex structures, there are no ground truth structures to compare with. Although the results from Figure 5 can provide us some hints (or intuition) to guess the reliability of AlphaFold 2 prediction, this guess still carries with some human bias. Therefore, an open question remains: how can we evaluate the reliability of AlphaFold 2 for the unknown complex structures? It should be noted that Although AlphaFold 2 can provide confidence values of the prediction, these confidence values are focused on local region (each protein) and do not consider the interactions between proteins. Therefore, AlphaFold-Eva, a deep learning-based method, is trained to solve this problem and to evaluate the confidence between protein interaction.

The top 4 features from feature selection (surface ratio, contact surface, dimension ratio, and total surface) are chosen as input features. The target feature is AlphaFold 2 prediction results. 4 hidden layers are selected, and each hidden layer contains 30 neurons, as shown in Figure 6 (a). It should be noted that the performance of the neural network is enhanced by adding more hidden layers and hidden neurons, but a more complicated network structure can also converge to an unintended local optimum. Therefore, overfitting or underfitting is another concern to disturb the optimization process (Cao et al. 2016; C. Xu et al. 2019; Xiong et al. 2020).

**Figure 6.**
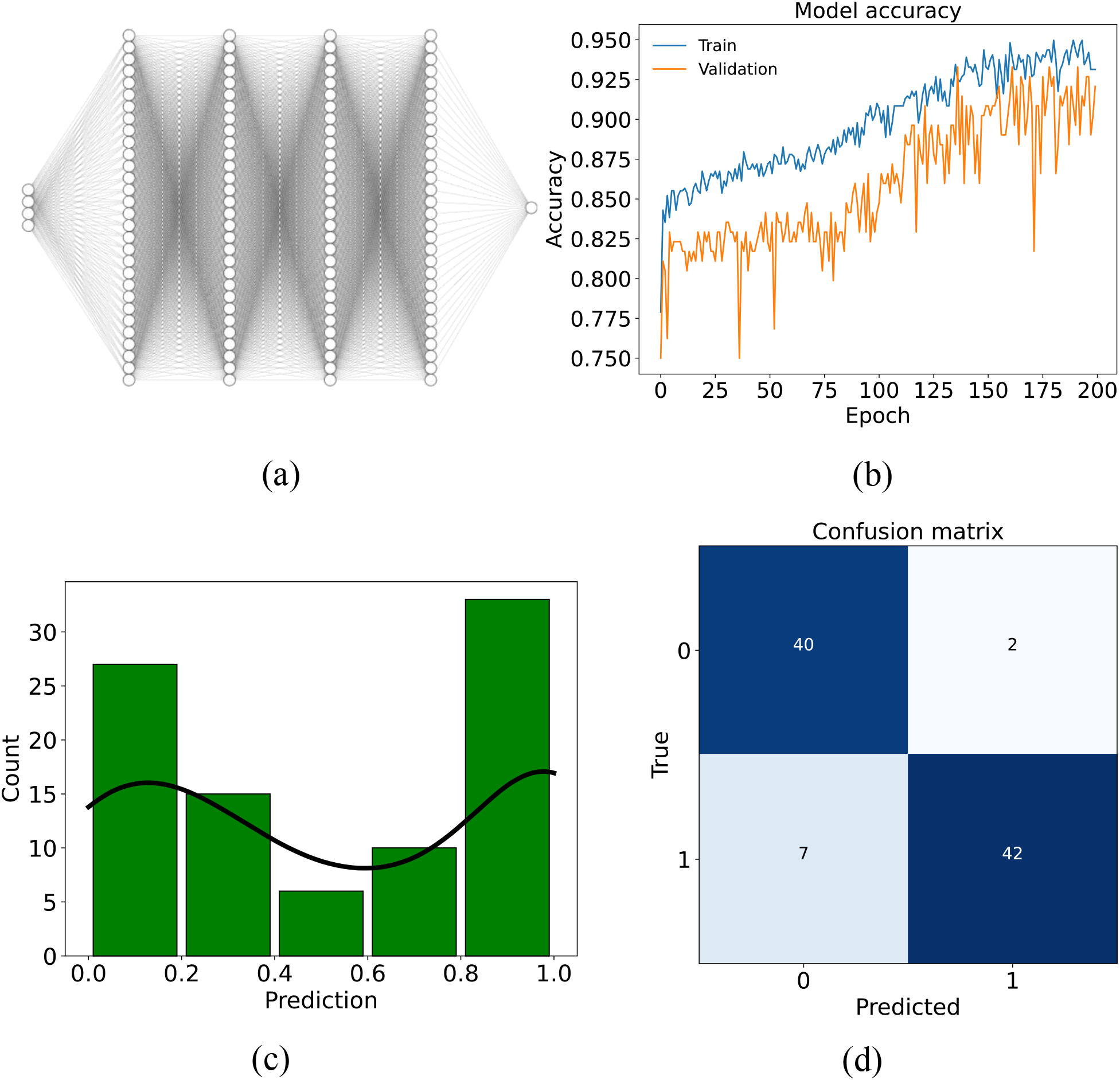
Neural network construction and application. (a) Neural network architecture; (b) The accuracy of models in train and validation dataset; (c) The distribution of prediction results of AlphaFold- Eva ranging from 0 to 1; (d) The confusion matrix for the test dataset.

To testify the performance of our neural network architecture and avoid overfitting or underfitting problems, initially, the whole input features are scaled using MinMaxScaler, and a box plot is used to eliminate the outliers. 90% of the data is used as training data and the rest is as test data. Next, the training data is split again and 80% of the data is treated as train data and the rest is as validation data where cross-validation is used to determine the best split (Sen et al. 2021).

RELU is used as the nonlinear activation for the first four layers (Zoph et al. 2018) and Sigmoid is for the last layer (Nwankpa et al. 2018), and the optimizer is Adam (Kingma et al. 2015). Figure 6 (b) shows the accuracy of the model in train and validation data. After 200 epochs, the accuracy of the trained AlphaFold-Eva can reach 95%. Meanwhile, the small deviation between train and validation after 200 epochs indicates that our trained model does not have the overfitting or underfitting problems and is a good fit model that suitably learns the train data and generalizes well to the validation data.

Next, we use the well-trained AlphaFold-Eva to predict test data. Figure 6 (c) shows the distribution of prediction values ranging from 0 to 1. The larger the value is, the less accurate prediction will be. For example, when the prediction value is 0.6, it means we have 60% confidence to confirm that the AlphaFold 2 prediction is bad. To compare the prediction values with true values from test data, we round the prediction values (the threshold value is 0.5). Figure 6 (d) is the confusion matrix where the diagonal shows the accurately predicted numbers via AlphaFold-Eva. The predicted accuracy is about 90%, indicating AlphaFold-Eva can accurately estimate the reliability of the AlphaFold 2.

### Application of AlphaFold-Eva

After validating the accuracy of AlphaFold-Eva, we use it to evaluate the reliability of AlphaFold 2 on unknown complex structures (the third dataset: Central Pair Complexes and recently released PDB files). Table 1 shows the input features after MinMaxScaler and predicted values using AlphaFold-Eva (the less the better). It shows that for complexes DYP30_DYP30 and 7KD9, the AlphaFold-Eva prediction values are 0.103764 and 0.106599, indicating we have about 90% confidence to tell that the prediction from AlphaFold 2 is correct but for 7CP2, we only have 36% confidence. For the rest of the complexes, we have almost 100% confidence to tell that the prediction from AlphaFold 2 is inaccurate. Then we compare the prediction structures from AlphaFold 2 with our unpublished Central Pair structures and recently released ground truth, as shown in Figure 7. It shows the predictions of DYP30_DYP30, 7KD9, and 7CP2 from AlphaFold 2 are reliable (green labels in Figure 7) but the rest of the predictions are not reliable.

**Table 1.** Input features after MinMaxScaler and predicted results using AlphaFold-Eva.

| Complex | Total surface | Contact surface | Surface ratio | Dimension ratio | Prediction |
| --- | --- | --- | --- | --- | --- |
| DYP30_DYP30 | 0.023964796 | 0.115880619 | 0.33570883 | 0.253203213 | 0.103764 |
| FAP101_CAM1 | 0.263603408 | 0.064296242 | 0.051385824 | 0.09420501 | 0.999852 |
| FAP114_FAP119 | 0.15174901 | 0.231208548 | 0.208122828 | 0.463829083 | 1.000000 |
| FAP194_FAP108 | 0.302731893 | 0.0449521 | 0.036616891 | 0.021953869 | 0.999995 |
| FAP227_FAP114 | 0.101455639 | 0.06779661 | 0.116295104 | 0.449597769 | 1.000000 |
| FAP227_FAP119 | 0.101642499 | 0.021002211 | 0.069856703 | 0.298534802 | 1.000000 |
| FAP305_FAP119 | 0.18510352 | 0.103721444 | 0.096035013 | 0.522302893 | 1.000000 |
| FAP360_FAP108 | 0.335245534 | 0.452652911 | 0.193733844 | 0.193207555 | 1.000000 |
| FAP16_FAP69 | 0.649357202 | 0.007921887 | 0.006909444 | 0.07760427 | 1.000000 |
| FAP16_FAP305 | 0.256016892 | 0.043478261 | 0.042510359 | 0.857890948 | 1.000000 |
| FAP194_FAP69 | 0.656831602 | 0.270449521 | 0.0634047 | 0.068971884 | 1.000000 |
| 7KD9 | 0.132465057 | 0.336588062 | 0.317426222 | 0.018256622 | 0.106599 |
| 7CP2 | 0.045836759 | 0.087324982 | 0.215840736 | 0.0224282 | 0.639445 |
| 6WTY | 0.16538979 | 0.024871039 | 0.050090404 | 0.116900948 | 0.999976 |
| 6XWP | 0.243870992 | 0.154937362 | 0.102019194 | 0.13875033 | 0.999963 |
| 6WZ1 | 0.068586965 | 0.049926308 | 0.12714929 | 0.041379082 | 0.962816 |
| 7LCV | 0.112770013 | 0.09137804 | 0.129639304 | 0.261042452 | 1.000000 |

**Figure 7.**
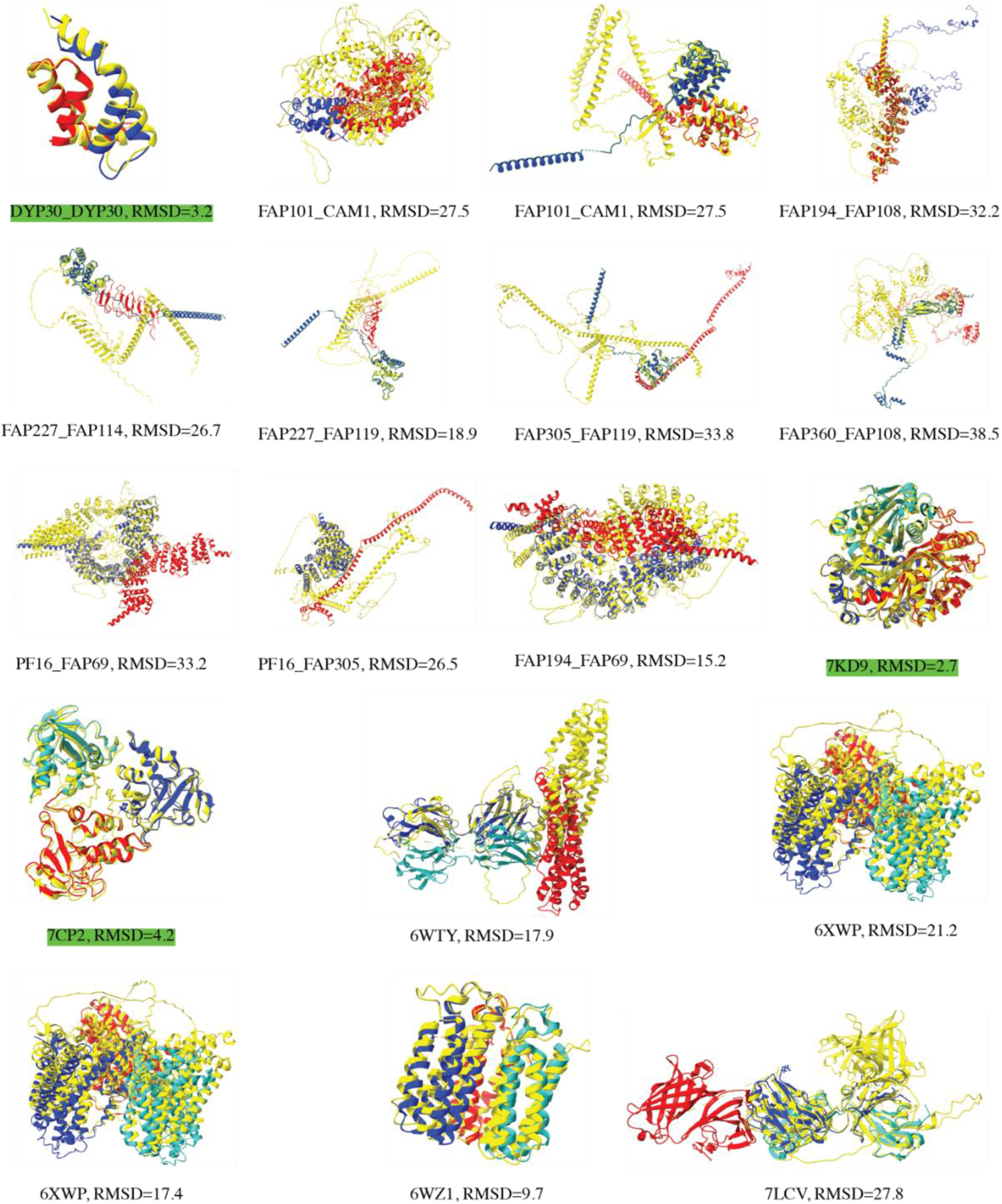
Comparison of AlphaFold 2 prediction structures with our unpublished central pair structures and recently released PDB data. The green label means AlphaFold 2 can provide a good prediction. Color code follows Figure 5.

Except for the 7CP2 complex, this observation is consistent with the conclusion we obtain from AlphaFold-Eva, demonstrating AlphaFold-Eva can provide a high accuracy when evaluating the AlphaFold 2 reliability on unknown complex structures.

## Further Discussion

From now on, we have tested the accuracy of AlphaFold 2 on complexes, but a question still remains: how about the performance of AlphaFold 2 on complexes if one of the components is missing? To test this thought, we have manually ignored one sequence of amino acids and put these incomplete sequences into AlphaFold 2, and part of the results (trimer) is shown in Figure 8, where left three panels are the comparison of AlphaFold 2 prediction (3 components) with the references and right three panels are the same structure comparison of AlphaFold 2 prediction with the references but with only 2 components. Figure 8 (a, left one) indicates that AlphaFold 2 cannot predict the protein interaction between protein 1 and proteins 2-3 accurately. This is consistent with our trained AlphaFold-Eva where the input features are [0.29058599, 0.15070007, 0.08551379, 0.22395017] and prediction value is 0.99997103. However, if we only consider the interaction between proteins 2 and 3, we can see that the AlphaFold 2 prediction matches the reference pretty well, which also agrees with AlphaFold-Eva where the input features are [0.12458891, 0.28997789, 0.29164829, 0.08437477] and prediction value is 0.14708968.

**Figure 8.**
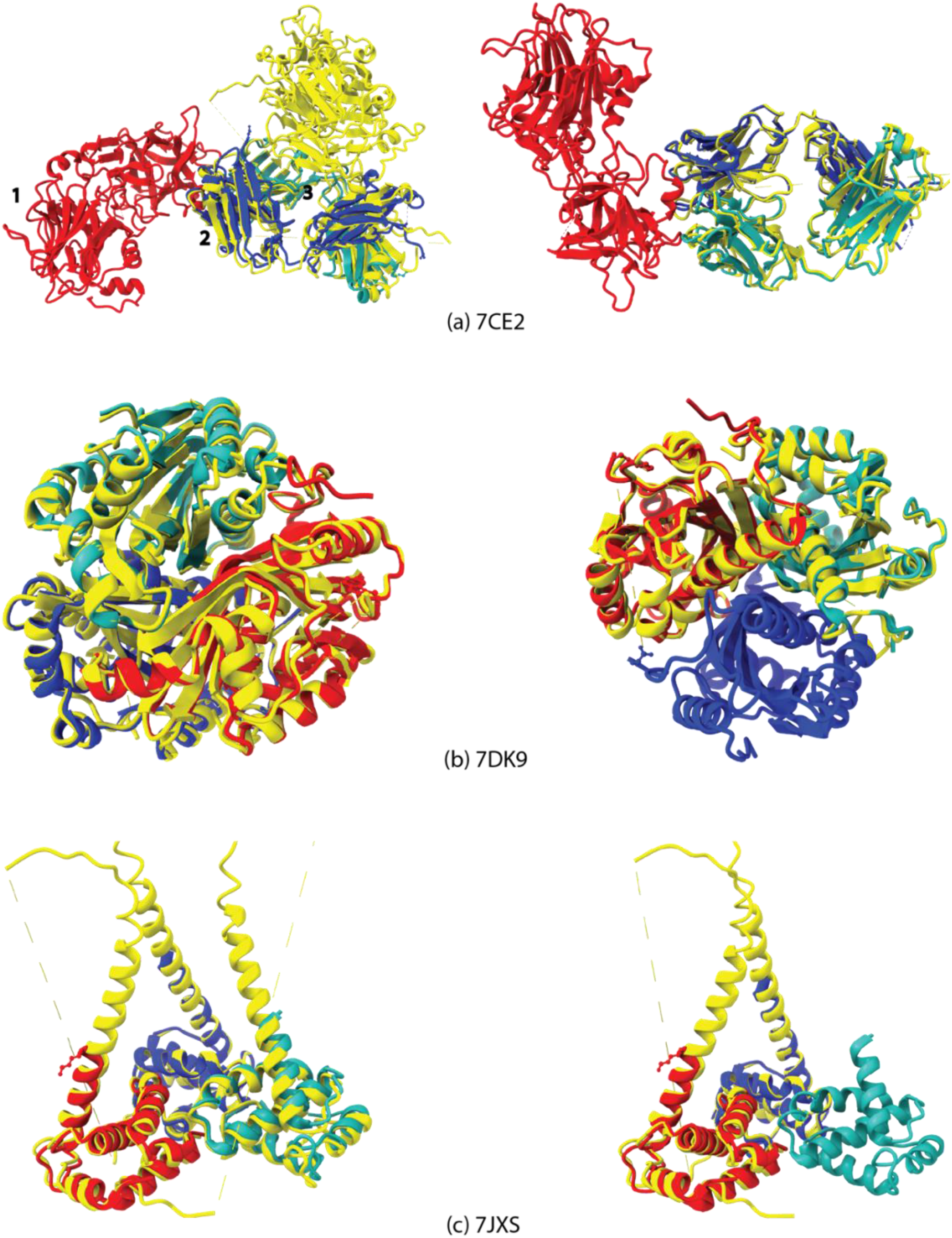
Comparison of AlphaFold 2 prediction structures with recently released PDB data. (a-c) Left, the whole sequences are considered; Right: one sequence of the complex is ignored. Color code follows Figure 5.

Next, we ignore the sequence of protein 1 (red) and only use the sequences of proteins 2 and 3, and the result is shown in Figure 8 (a, right one), which shows a similar result with Figure 8 (a, left one) of subcomplex (proteins 2 and 3). This result indicates the existence of protein 1 does not influence performance of AlphaFold 2. Figure 8 (b and c) shares the similar phenomenon with Figure 8 (a). Note that all the tested complexes are not included in AlphaFold 2 database and the interactions among each protein (domain) are not twisted.

Therefore, through these examples, we can safely refer that when predicting a complicated complex, we can use AlphaFold 2 to predict some sub-complexes of it. During this process, AlphaFold-Eva can evaluate the reliability of each prediction and obtain the confidence value of each sub-complex, and finally we can determine which part of AlphaFold 2 prediction is trustworthy.

## Conclusion

In this study, protein-protein complexes are used to testify the accuracy of AlphaFold 2. We show that AlphaFold 2 has the potential ability to predict the protein-protein complex structures where the performance for homomer (61.7%) is generally higher than that for heteromer (48.8%). This is because in homomer the contact surface is usually larger than that in heteromer, demonstrating that AlphaFold 2 can learn more information from the homomer complex interfaces. Moreover, structure dimension ratio is another important factor to influence the accuracy of AlphaFold 2, showing that the more cubic the structure is, the higher the accuracy will be. Finally, AlphaFold-Eva is trained to evaluate the reliability of AlphaFold 2 on unknown complex prediction where the accuracy of our trained model is 90%. As a limitation, AlphaFold-Eva is only suitable for the default AlphaFold 2, not applicable for the will-be-released AlphaFold-Multimer (Evans et al. 2021). However, part of the works can be the complementary explanation for Evans et al.’s (2021) results. Meanwhile, our work such as feature-importance selection will provide some clues for the DeepMind to improve their AlphaFold-Multimer (Evans et al. 2021).

The source code and weights for AlphaFold-Eva are available online (https://github.com/xiong19912010/AlphaFold-Eva).

## Acknowledgment

The computing for this project was performed at the FARNAM Supercomputing Center at Yale University.

